# Clustering and rapid long-range motility behavior of bacteria by type IV pili

**DOI:** 10.1101/525790

**Authors:** Chinedu S. Madukoma, Nydia Morales Soto, Anne E. Mattingly, Shaun W. Lee, Joshua D. Shrout

## Abstract

Many organisms coordinate to move and colonize over surfaces. Bacteria such as *Pseudomonas aeruginosa* exhibit such surface motility as a precursor step to forming biofilms. Here we show a group surface motility where small groups of *P. aeruginosa* use their type IV pili (TFP) appendages over long-distances. Small cell clusters employ their TFP to move multiple cell lengths in fractions of a second and form new multicellular groups. Given the length scale and speed of displacement, cells appear to “snap” to a new position and then resume their previous behavior. The same long range TFP action also leads to rapid community contraction of sparsely arranged cell clusters. Cluster development and snapping motility does not require exogenous DNA or extracellular polysaccharides.

## Introduction

TFP function as motility appendages by an extension, tip attachment, and retraction order of action [1]. These thin structures (~6 nm diameter) range from 0.5-7.0 μm in length and can generate pulling forces exceeding 100 pN [1–5]. For the bacterium *Pseudomonas aeruginosa*, TFP are known to confer motility phenotypes known as twitching and walking. *P. aeruginosa* also employs its polar flagellum to confer motility modes called swimming and swarming. The transition between flagellar-mediated motility to TFP-mediated motility is not well understood. For example, while swarming requires active flagella, the absence or over expression of TFP have been shown to influence the overall swarm phenotype [6]. Here we show that TFP facilitate a cluster development and rapid motility phenotype in a subset of cells following flagellar-driven swarming community expansion. We find that *P. aeruginosa* TFP can extend multiple cell lengths (>30μm) and then retract to translocate small clusters of cells to join with other cell clusters in less than one second.

## Results and Discussion

There are two modes of *P. aeruginosa* surface motility that are routinely investigated using agar motility plate assays. Flagellar-mediated swarming is studied using 0.4%-0.6% agar and TFP-mediated twitching is studied using 1.0% agar. Both swarming and twitching are commonly assessed in these plate assays as growing and expanding *P. aeruginosa* populations that develop over hours to days. Here we detail surface motility behavior that occurs on surfaces between those conditions typically used to examine swarming and twitching (0.4-1.0% agar). We find that as *P. aeruginosa* colonies expand from the point of inoculation, small clusters of cells advance beyond the larger group. Using this approach, an open-air plate was required—covering the agar (with a glass coverslip) does not promote the same cluster development. We find this clustering phenotype is conserved among *P. aeruginosa* strains, as all common laboratory strains tested (PA14, PAO1, MPAO1, PAK and PAO1C) develop clusters to some degree (Figure S1).

Inspection of these clusters over time shows they are transient and facilitate the restructuring of the community. Cells cluster, divide into smaller groups, and recluster numerous times within a minute. Some of these clusters move multiple cell lengths to join other clusters or the larger advancing colony within less than a second (Figure 1—top panels), thus appearing to “snap” from one region of the frame to another (Movie S1).

**Fig. 1.**
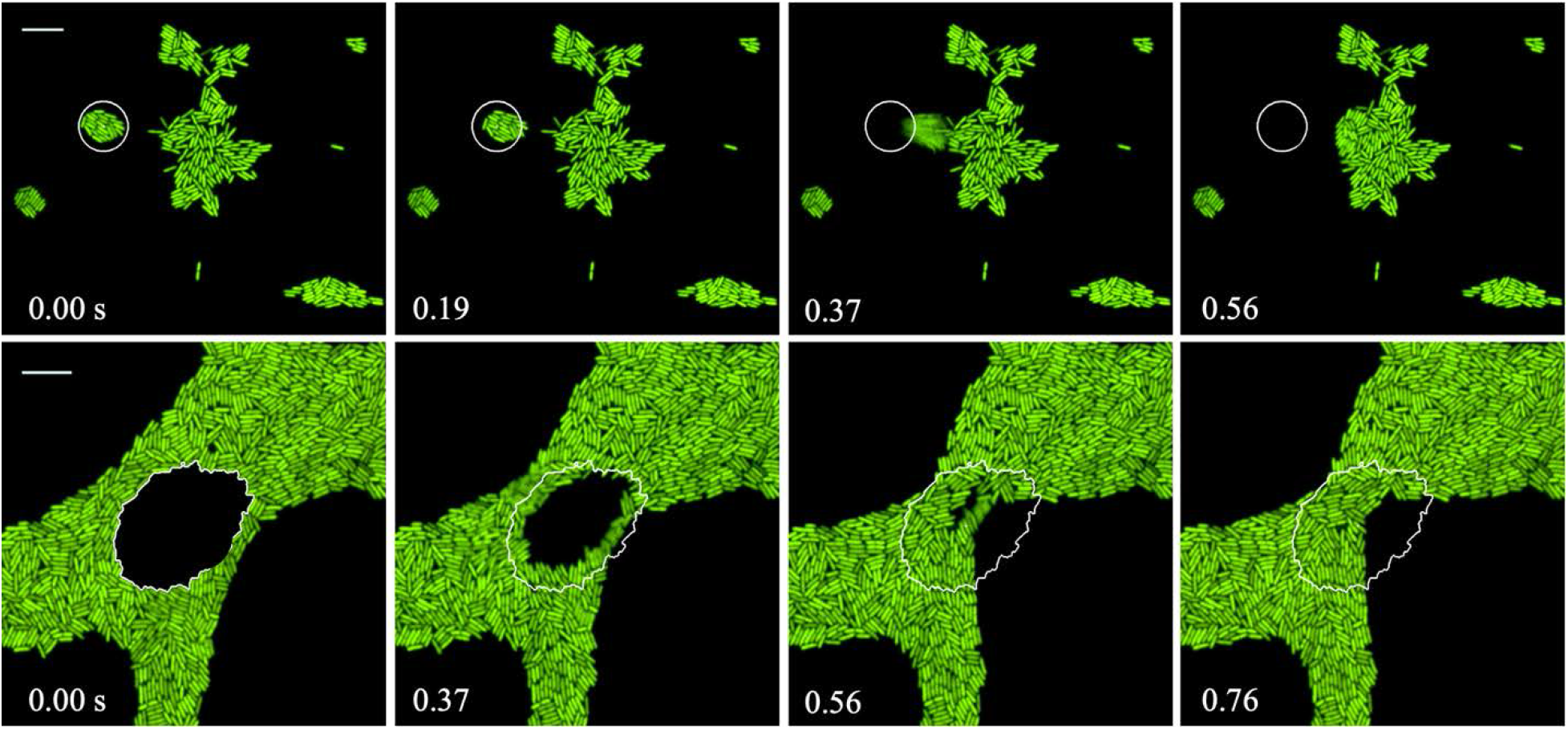
*P. aeruginosa* snapping and community contraction in less than 1 second. Time-lapse sequences of representative *P. aeruginosa* snapping and community contraction events are shown. Top row: advancing cluster (identified with white circle at the 0 s) prepares to snap by enhacing cluster arrangement (0.19 s), begins moving (0.37 s), and joins another cluster (0.56 s). Bottom row: A contraction snapping event. The open area surrounded by cells (traced in white— spanning 469.87 μm^2^) is covered within 0.76 s. Scale bar represents 10 μm.

In all, we track cells in 115 of these snapping clusters in various assays and specifically quantify 70 snapping events in experiments containing 0.8% agar (Table 1). These snapping clusters contain an average of 17.5 cells and travel a maximum distance of 17.8 μm, or ~4 cell lengths. Cluster size does not correlate with distance traveled or snapping duration. Likewise, no apparent trend changes are attributable to the original time of assay inoculation. However, the total distance traveled by snapping clusters strongly relates with water availability in these surface motility assays. The maximum snapping distance on 0.45% agar assays is 37 μm or ~9 cell lengths, while the maximum distance cells snap on 1.0% agar is 10 μm.

**Table 1.**
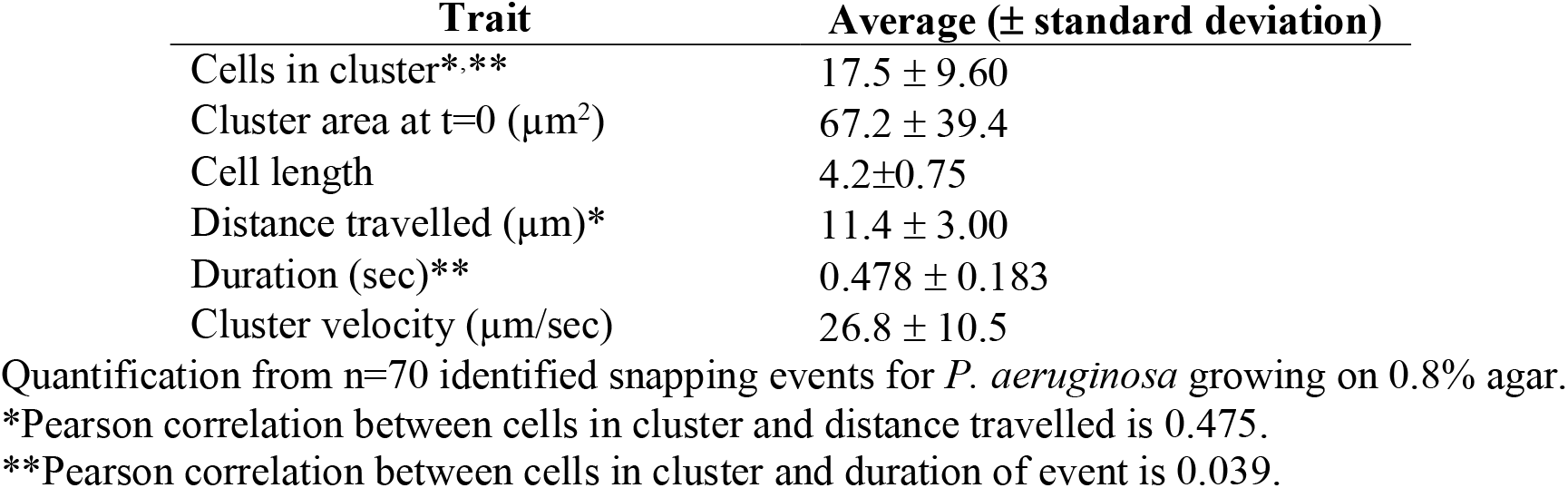
*P. aeruginosa* snapping behavior dynamic traits.

| Trait | Average (± standard deviation) |
| --- | --- |
| Cells in cluster*** | $17.5 \pm 9.60$ |
| Cluster area at t=0 ($\mu\text{m}^2$) | $67.2 \pm 39.4$ |
| Cell length | $4.2 \pm 0.75$ |
| Distance travelled ($\mu\text{m}$)* | $11.4 \pm 3.00$ |
| Duration (sec)** | $0.478 \pm 0.183$ |
| Cluster velocity ($\mu\text{m}/\text{sec}$) | $26.8 \pm 10.5$ |

Quantification from n=70 identified snapping events for *P. aeruginosa* growing on 0.8% agar.
\*Pearson correlation between cells in cluster and distance travelled is 0.475.
\*\*Pearson correlation between cells in cluster and duration of event is 0.039.

Larger less ordered groups also exhibit this long-distance rapid cell movement as a means of community contraction. A representative example is shown in Figure 1 (bottom panels) where an open area surrounded by cells (traced in white—spanning 469.87 μm^2^) collapses within 0.76 seconds.

Snapping behavior requires *P. aeruginosa* TFP. Isogenic mutants deficient for TFP fail to form the small aggregates and no sub-second rapid behavior is detected (Figure 2). Further, functional TFP retraction is required as the retraction-deficient *pilU* strain also does not form clusters and exhibits behavior like the *pilA* strain. When one considers the size of PilA monomers [7] and TFP assembly [8] required to mediate snapping, nearly 17,000 PilA monomers are required to assemble TFP capable of the average event measured in our experiments. Accordingly, the pilus required to mediate the maximum distance event (of 37 μm) contains a staggering 35,200 PilA monomers.

**Figure 2.**
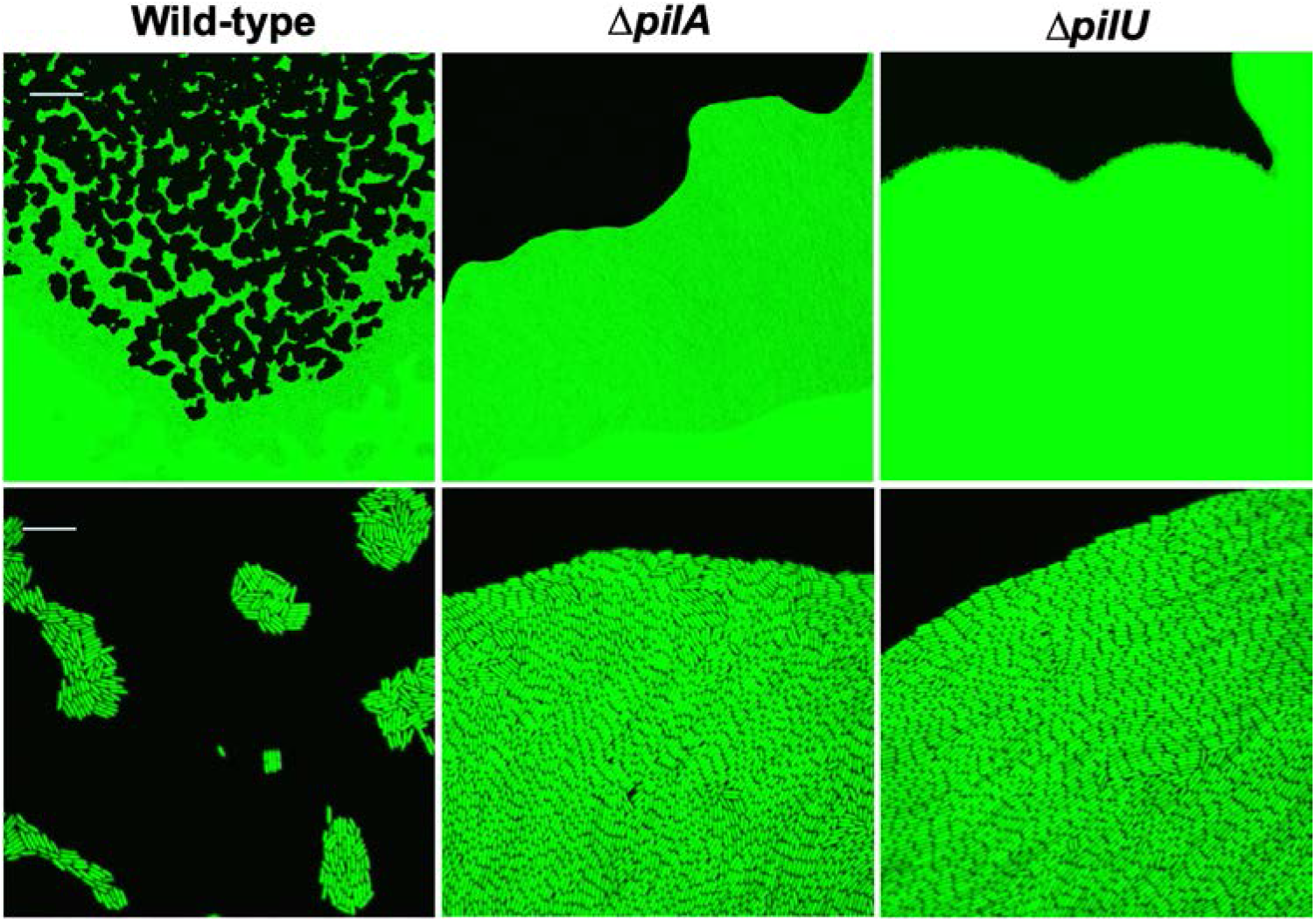
Advancing colony phenotype of *P. aeruginosa* on motility assay plates containing 0.8% agar. Top (10× magnification) demonstrates cluster development by wild-type beyond the advancing swarm zone. TFP-deficient (Δ*pilA*) and TFP retraction-deficient (Δ*pilU*) strains make no clusters. Bottom row (100× magnification) shows arrangements of single cells, and Δ*pilA* and Δ*pilU* appear highly ordered and tightly packed. Scale bars represent 100 *μ*m (top) and 10 μm (bottom).

A lag time is seen after plate assay inoculation before snapping is observed. We also see that a functional flagellum is a required precursor to this snapping motility phenotype. On 0.5% agar assays that promote robust swarming, the lag time for cluster development and snapping is 1.5-2 days (as the plates dry out). However, using an agar range of 0.8%-1.0% promotes cluster formation and snapping within a few hours post inoculation. Cells missing their polar flagellum (*fliC*) are widely dispersed and fail to develop the small clusters that form beyond the advancing large population on these surface assays (Figure 3). Thus, it appears that flagellar-mediated swarming can serve as an initial step to establish the needed population of cells to subsequently form cell clusters that will snap together using their TFP.

**Figure 3.**
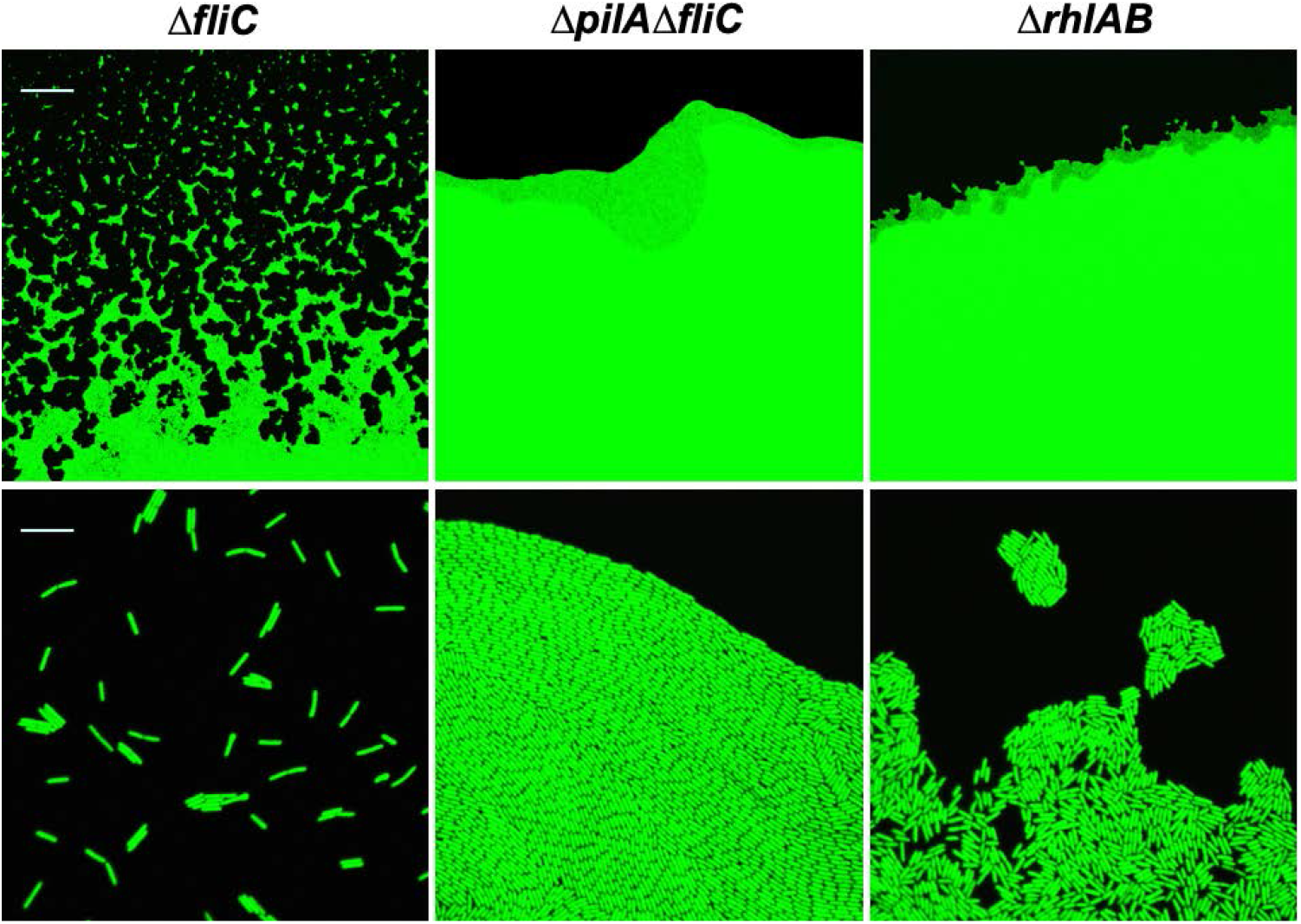
Colony edge and clustering phenotype of *P. aeruginosa* mutants on 0.8% agar. Top (10× magnification): Flagella-deficient (Δ*fliC*) are mostly sparsely distributed at the single-cell level, while the appendage-deficient strain (Δ*pilA*Δ*fliC*) do not form cells clusters. Rhamnolipid-deficient (Δ*rhlAB*) cells make a reduced number of smaller clusters closer to the colony edge. Bottom row (100× magnification) show arrangements of single cells (mono-layer). Images were obtained using confocal microscopy twelve hours after inoculation. Top (10× magnification) Scale bars represent 100 μm (top) and 10 μm (bottom).

Having established TFP as the appendages that mediate snapping, we investigate the additional biological factors required. We find that snapping also requires the surfactant rhamnolipid. Cells from a rhamnolipid-deficient strain (Δ*rhlAB*) can move away from the uniformly expanding *P. aeruginosa* colony and form some clusters, but these bacteria never exhibit snapping motion (Figure 3).

We next examine the potential role of DNA as a marker and promoter of snapping motility as previous studies have shown the ability of *P. aeruginosa* TFP to track to exogenous DNA as a component of biofilm development [9,10]. We find that snapping precedes the presence of detectable DNA in these small cell groups. Snapping is observed to occur in the same experiment as “explosive cell lysis” [11]. Figure 4 shows snapping of a small cluster in a region with no exogenous DNA while a cell simultaneously explodes to release DNA in a separate region (sequence included as Movie S2). Thus, we conclude that snapping does not require TFP-DNA interaction and is distinct from TFP actions mediated by exogenous DNA.

**Figure 4.**
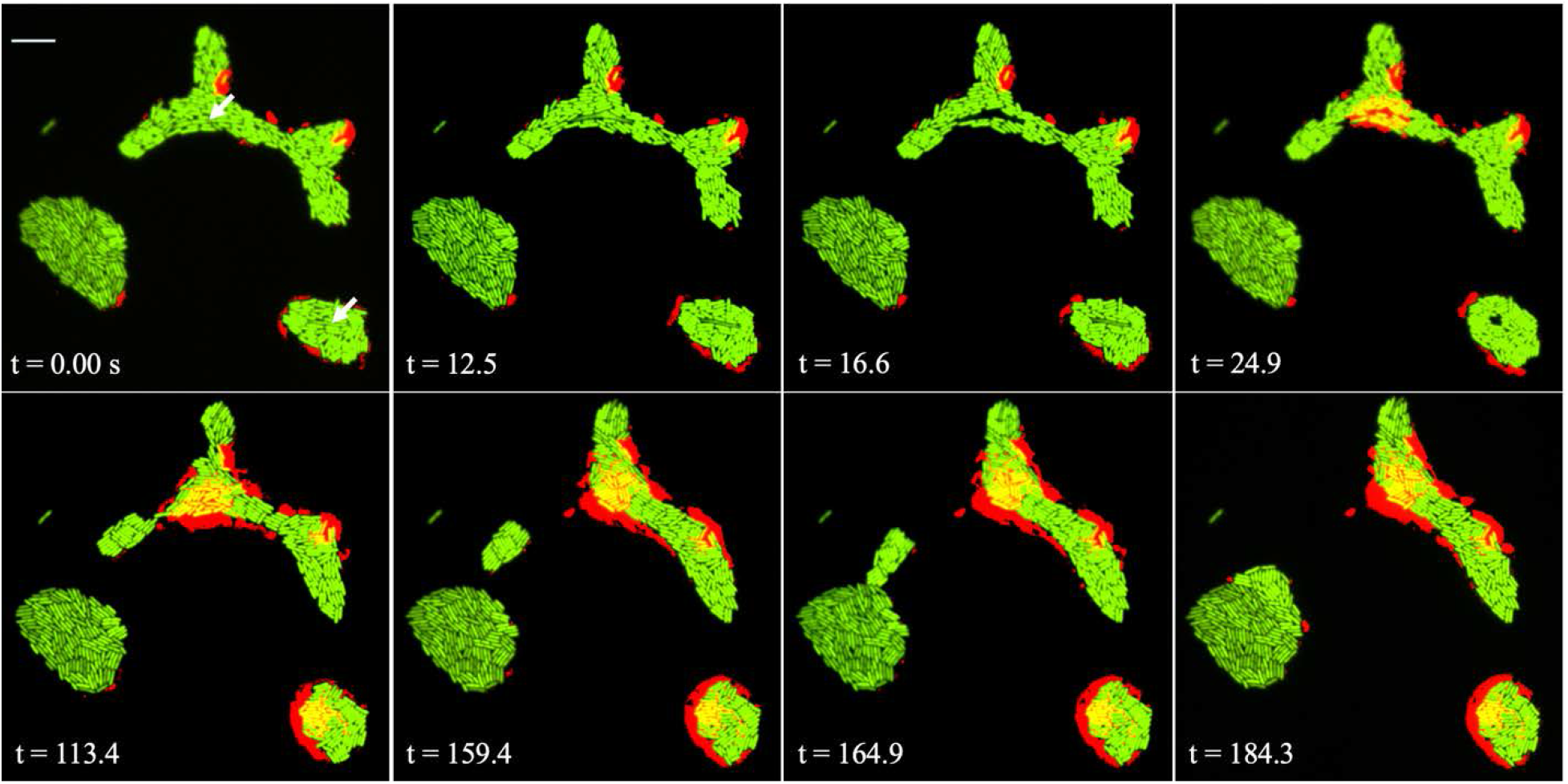
Sequence of events demonstrating eDNA release from a lysing cell within the same time span as snapping motility that occurs elsewhere in the same frame. A cell ready to lyse (top arrow, 0.00 s) ceases production of GFP (12.5 s), explodes by 16.6 s, and releases its exogenous DNA at 24.9 s. The red color indicates the DNA released by the lysing cell. A cluster begins to pull away (113.4 s) from the cluster exhibiting a high concentration of high eDNA and snaps onto a different cluster by the 116.9 s. Scale bar represents 10 μm.

Since DNA did not serve as an attractant for snapping clusters, we next investigate the *P. aeruginosa* polysaccharides Pel and Psl, which have been shown important to *P. aeruginosa* surface motility and aggregation as a stage in biofilm development [12,13]. Similarly, the bacterium *Myxococcus xanthus* is well known to require exopolysaccharide to confer “social” TFP-mediated motility [14–16]. However, we find that the Pel and Psl polysaccharides for *P. aeruginosa* do not confer cluster development or the snapping phenotype. Elimination of either or both the Pel and Psl polysaccharides actually leads to increased cluster development and equivalent snapping events to that we observe for the wildtype (Figure 5).

**Figure 5.**
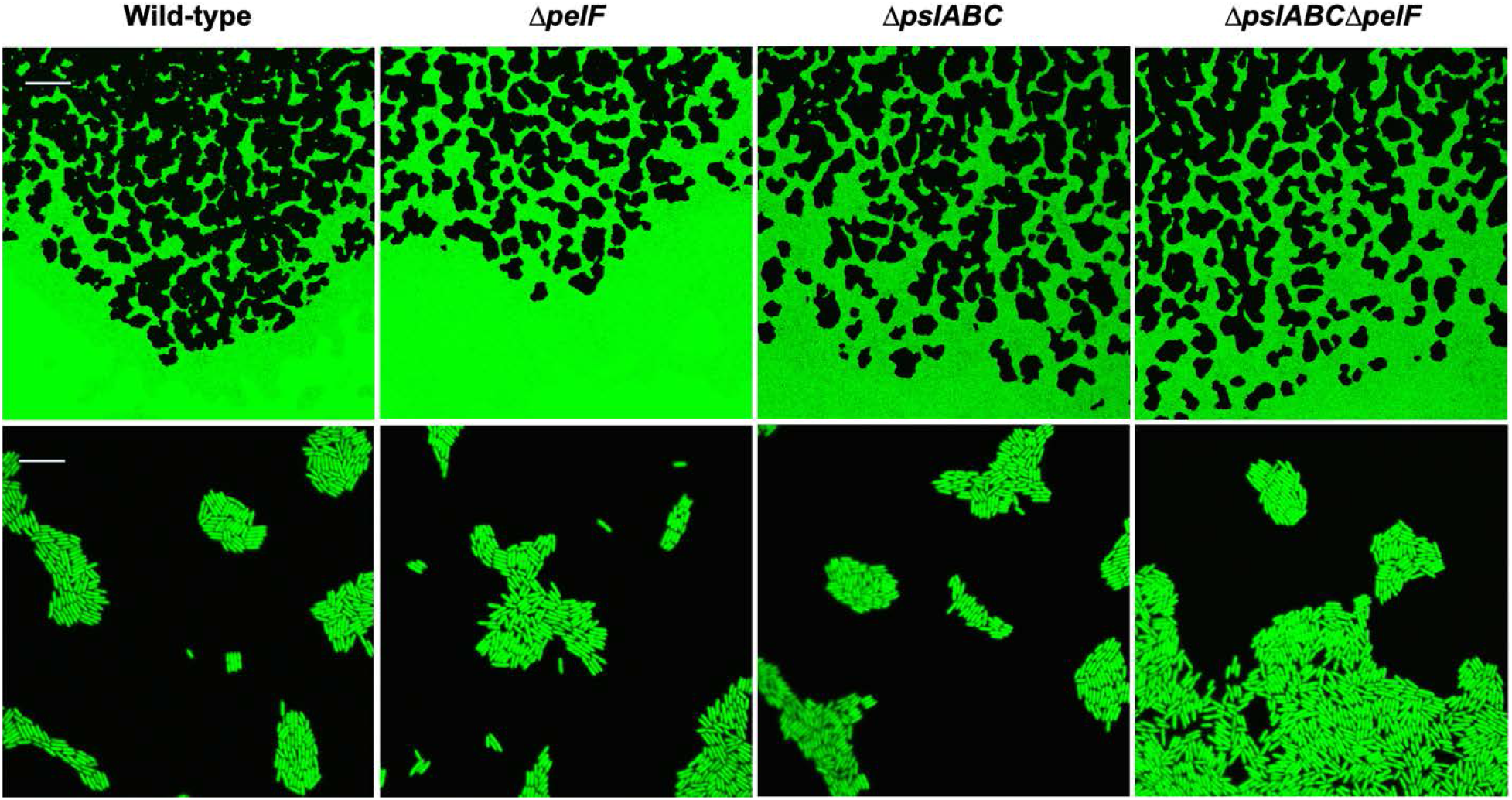
Top (10× magnification) demonstrates cluster formation by Pel-deficient (Δ*pelF*), Psl-deficient (Δ*pslABC*), and polysaccharide-deficient (Δ*pslABC*Δ*pelF*) in similar fashion as the wild-type. Bottom row (100× magnification) shows the arrangements of single cells (mono-layer) within these clusters and advancing swarm edges. Bottom row (100× magnification) show clusters at of single cell level. Scale bar represent 100 *μ*m (top) and 10 *μ*m (bottom).

Here we detail TFP-dependent snapping in the bacterium *P. aeruginosa*. Our probing of cluster development on semi-solid surfaces suggests that snapping occurs readily as small groups of *P. aeruginosa* cells rapidly restructure the surrounding community within seconds.

## Materials and Methods

### Bacteria strains and growth medium

Strains of *Pseudomonas aeruginosa* used in this study are included in Table S1. Cultures were streaked from frozen (−80°C) stocks onto LB agar plates (1.5% wt/vol) and incubated at 37°C overnight. Isolated colonies selected and grown planktonically in 6 mL FAB with 30 mM glucose [6,17] at 37°C with shaking at 240 rpm for 12-15 hrs.

### Isogenic mutant construction

Primers and gBLOCK nucleotides used for this work are included in Table S2. Plasmid pCSM1 was constructed to allow creation of a markerless Δ*pslABC* mutant. pCSM1 was constructed by inserting a single gBLOCK of double-stranded DNA (IDT, Coralville, IA) containing both regions upstream of the *pslA* gene and downstream of the *pslC* gene according to the *Pseudomonas* genome database [18]; this gBLOCK was restricted using EcoRI and HindIII and ligated into the pEX18Ap vector between the EcoRI and HindIII sites and expressed in *E. coli* DH5α.

Similarly, plasmids pNMS and pCSM103 were constructed to allow creation of a markerless Δ*rhlAB* and Δ*fliC* mutants. pNMS utilized a single gBLOCK containing both regions upstream of the *rhlA* gene and downstream of the *rhlB* gene and pCSM103 contained regions upstream and downstream of the *fliC* gene. These amplified gBLOCK products were ligated into the pEX18Gm plasmid [19] following SLiCE protocols [20].

Mutations were introduced using conjugational mating. Single cross-over recombinants were selected on LB plates (1.5% agar wt/vol), augmented with 100μg/ml carbenicillin (pNMS, pCSM1 and pCSM103) or 100μg/ml gentamycin (pEX18GmΔ*pelF*). Double crossover recombinants were then selected on 5% wt/vol sucrose LB plates. Deletions were confirmed by comparing PCR amplification of the target region with the PAO1C wildtype.

### Constitutive fluorescent strain construction

Plasmids harboring a Tn7 region containing green fluorescent protein under control of a constitutive strong promoter were incorporated into *P. aeruginosa* strains by conjugational mating. Either plasmid pBK miniTn7-gfp2 (harboring gentamycin resistance) or pBK miniTn7-*gfp3* (harboring kanamycin and streptomycin resistance) were incorporated adjacent to *glmS* on the *P. aeruginosa* chromosome using helper plasmid pUX-BF13 and mobilization plasmid pRK600. Transgenic cells were selected on LB agar supplemented with 100μg/mL gentamycin or 250μg/mL kanamycin and 250μg/mL streptomycin and confirmed by PCR detection of the chromosomal insertions and microscopy inspection for fluorescence.

### Surface motility assays

Surface motility plate assays were composed of modified fastidious anaerobe broth (FAB) media that included 0.1% casamino acid (wt/vol) and no ammonium sulfate ((NH_4_)_2_SO_4_) [6,17]. Surface motility was evaluated over a range of Noble Agar concentrations between 0.4%-3.0% (wt/vol). Plate assays were spot inoculated by pipetting 1μm log-phase cells onto the center of the 60mm Petri dish and quickly transferred to a laminar hood for 3-5 minutes with lids partially open, promoting rapid absorption of the inoculation droplet into the motility agar. All plates were inverted and incubated at 30°C in a humidity-controlled (85% RH) incubator for 15-30 h. Prior to imaging, motility plates were equilibrated to room temperature for 45-60 minutes.

### Microscopy and imaging

Imaging was carried out using a Nikon Eclipse Ti-E Inverted microscope equipped with Plan Fluor 10X DIC LN1 and 100X LU Plan Fluor objectives and Andor DU-888 X-10006 camera with Emission wavelength = 535, Excitation wavelength = 470.

### Image processing and analysis

Images were processed using Nikon NIS-Element software and further analyzed using Fiji image processing package (3). Using tools in these software packages, size of clusters, distance and speed travelled by a cluster were determined.

## Acknowledgements

This work was supported by NIH grants 1R21AI109417 (JDS) and 1DP2OD008468 (SWL).

The PA14Δ*pelF* strain and pEX18GmΔ*pelF* plasmid were provided by Matthew Parsek, University of Washington-Seattle.

## Supplementary Materials

**Table S1.**
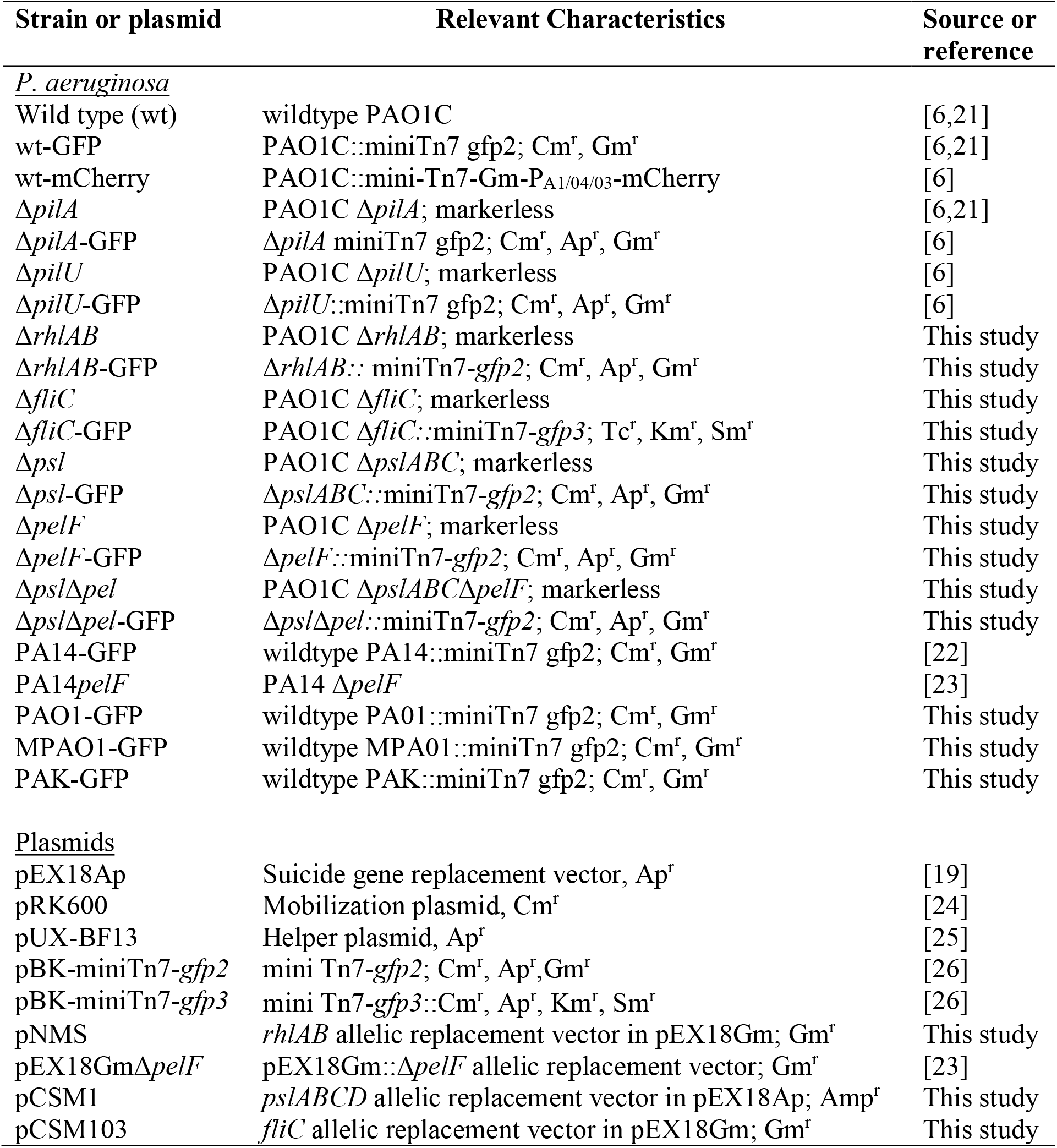
Strain list

**Table S2.**
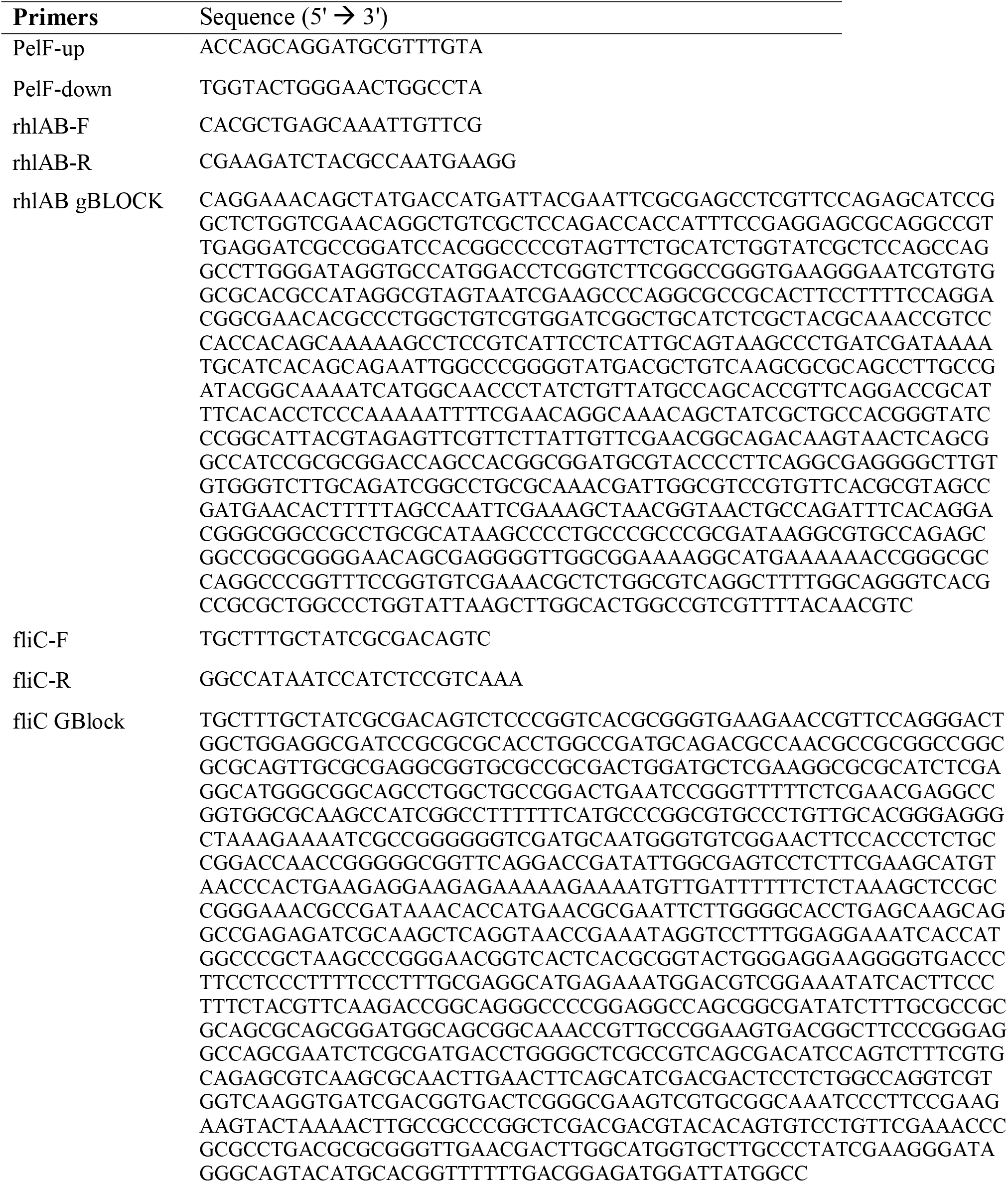
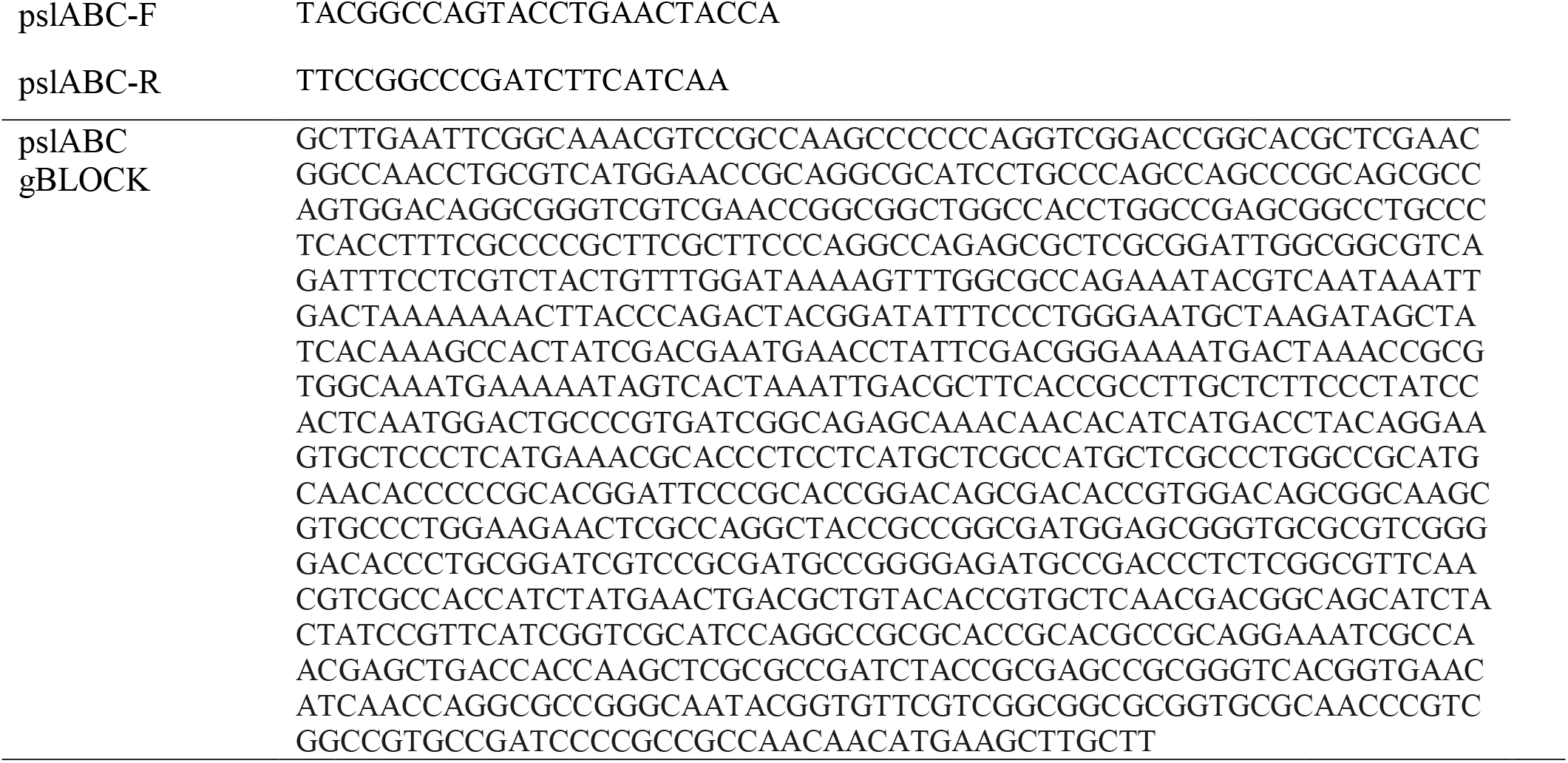
Primers and gBLOCK sequences

**Fig. S1.**
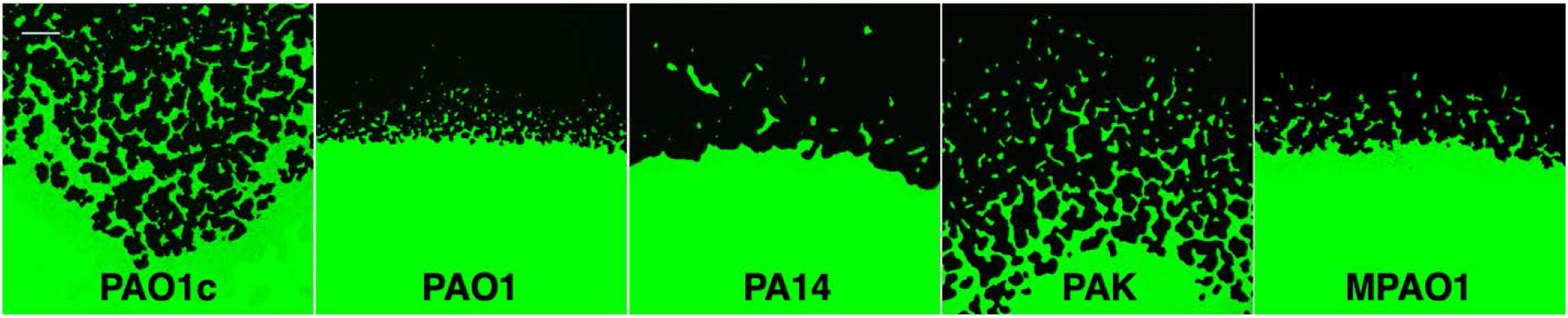
Cluster formation is conserved among wild-type *P. aeruginosa* strains on 0.8% agar. Scale bar represent 100 μm.

## Supplemental Movie Captions

Movie S1. *P. aeruginosa* snapping motility. The cluster in the upper right of frame undergoes some cell-cell rearrangement and then snaps to the larger cluster from right-to-left. (5.3 frames per second)

Movie S2. *P. aeruginosa* rapid community contraction. The open area surrounded by cells (spanning 469.87 μm^2^) is covered within 0.76 s.

