## Supplemental material for "Clustering and rapid long-range motility behavior of bacteria by type IV pili"

**Supplementary Materials:**

Tables S1-S2

Figure S1

Movies S1-S2

**Table S1. Strain list**

| **Strain or plasmid** | **Relevant Characteristics** | **Source or reference** |
|---|---|---|
| <u>*P. aeruginosa*</u> | | |
| Wild type (wt) | wildtype PAO1C | [6,21] |
| wt-GFP | PAO1C::miniTn7 gfp2; Cm$^r$, Gm$^r$ | [6,21] |
| wt-mCherry | PAO1C::mini-Tn7-Gm-P$_{A1/04/03}$-mCherry | [6] |
| Δ*pilA* | PAO1C Δ*pilA*; markerless | [6,21] |
| Δ*pilA*-GFP | Δ*pilA* miniTn7 gfp2; Cm$^r$, Ap$^r$, Gm$^r$ | [6] |
| Δ*pilU* | PAO1C Δ*pilU*; markerless | [6] |
| Δ*pilU*-GFP | Δ*pilU*::miniTn7 gfp2; Cm$^r$, Ap$^r$, Gm$^r$ | [6] |
| Δ*rhlAB* | PAO1C Δ*rhlAB*; markerless | This study |
| Δ*rhlAB*-GFP | Δ*rhlAB::* miniTn7-*gfp2*; Cm$^r$, Ap$^r$, Gm$^r$ | This study |
| Δ*fliC* | PAO1C Δ*fliC*; markerless | This study |
| Δ*fliC*-GFP | PAO1C Δ*fliC::*miniTn7-*gfp3*; Tc$^r$, Km$^r$, Sm$^r$ | This study |
| Δ*psl* | PAO1C Δ*pslABC*; markerless | This study |
| Δ*psl*-GFP | Δ*pslABC::*miniTn7-*gfp2*; Cm$^r$, Ap$^r$, Gm$^r$ | This study |
| Δ*pelF* | PAO1C Δ*pelF*; markerless | This study |
| Δ*pelF*-GFP | Δ*pelF::*miniTn7-*gfp2*; Cm$^r$, Ap$^r$, Gm$^r$ | This study |
| Δ*psl*Δ*pel* | PAO1C Δ*pslABC*Δ*pelF*; markerless | This study |
| Δ*psl*Δ*pel*-GFP | Δ*psl*Δ*pel::*miniTn7-*gfp2*; Cm$^r$, Ap$^r$, Gm$^r$ | This study |
| PA14-GFP | wildtype PA14::miniTn7 gfp2; Cm$^r$, Gm$^r$ | [22] |
| PA14*pelF* | PA14 Δ*pelF* | [23] |
| PAO1-GFP | wildtype PA01::miniTn7 gfp2; Cm$^r$, Gm$^r$ | This study |
| MPAO1-GFP | wildtype MPA01::miniTn7 gfp2; Cm$^r$, Gm$^r$ | This study |
| PAK-GFP | wildtype PAK::miniTn7 gfp2; Cm$^r$, Gm$^r$ | This study |
| <u>Plasmids</u> | | |
| pEX18Ap | Suicide gene replacement vector, Ap$^r$ | [19] |
| pRK600 | Mobilization plasmid, Cm$^r$ | [24] |
| pUX-BF13 | Helper plasmid, Ap$^r$ | [25] |
| pBK-miniTn7-*gfp2* | mini Tn7-*gfp2*; Cm$^r$, Ap$^r$,Gm$^r$ | [26] |
| pBK-miniTn7-*gfp3* | mini Tn7-*gfp3*::Cm$^r$, Ap$^r$, Km$^r$, Sm$^r$ | [26] |
| pNMS | *rhlAB* allelic replacement vector in pEX18Gm; Gm$^r$ | This study |
| pEX18GmΔ*pelF* | pEX18Gm::Δ*pelF* allelic replacement vector; Gm$^r$ | [23] |
| pCSM1 | *pslABCD* allelic replacement vector in pEX18Ap; Amp$^r$ | This study |
| pCSM103 | *fliC* allelic replacement vector in pEX18Gm; Gm$^r$ | This study |

**Table S2. Primers and gBLOCK sequences**

| **Primers** | Sequence (5' → 3') |
|---|---|
| PelF-up | ACCAGCAGGATGCGTTTGTA |
| PelF-down | TGGTACTGGGAACTGGCCTA |
| rhlAB-F | CACGCTGAGCAAATTGTTCG |
| rhlAB-R | CGAAGATCTACGCCAATGAAGG |
| rhlAB gBLOCK | CAGGAAACAGCTATGACCATGATTACGAATTCGCGAGCCTCGTTCCAGAGCATCCG<br>GCTCTGGTCGAACAGGCTGTCGCTCCAGACCACCATTTCCGAGGAGCGCAGGCCGT<br>TGAGGATCGCCGGATCCACGGCCCCGTAGTTCTGCATCTGGTATCGCTCCAGCCAG<br>GCCTTGGGATAGGTGCCATGGACCTCGGTCTTCGGCCGGGTGAAGGGAATCGTGTG<br>GCGCACGCCATAGGCGTAGTAATCGAAGCCCAGGCGCCGCACTTCCTTTTCCAGGA<br>CGGCGAACACGCCCTGGCTGTCGTGGATCGGCTGCATCTCGCTACGCAAACCGTCC<br>CACCACAGCAAAAAGCCTCCGTCATTCCTCATTGCAGTAAGCCCTGATCGATAAAA<br>TGCATCACAGCAGAATTGGCCCGGGGTATGACGCTGTCAAGCGCGCAGCCTTGCCG<br>ATACGGCAAAATCATGGCAACCCTATCTGTTATGCCAGCACCGTTCAGGACCGCAT<br>TTCACACCTCCCAAAAATTTTCGAACAGGCAAACAGCTATCGCTGCCACGGGTATC<br>CCGGCATTACGTAGAGTTCGTTCTTATTGTTCGAACGGCAGACAAGTAACTCAGCG<br>GCCATCCGCGCGGACCAGCCACGGCGGATGCGTACCCCTTCAGGCGAGGGGCTTGT<br>GTGGGTCTTGCAGATCGGCCTGCGCAAACGATTGGCGTCCGTGTTCACGCGTAGCC<br>GATGAACACTTTTTAGCCAATTCGAAAGCTAACGGTAACTGCCAGATTTCACAGGA<br>CGGGCGGCCGCCTGCGCATAAGCCCCTGCCCGCCCGCGATAAGGCGTGCCAGAGC<br>GGCCGGCGGGGAACAGCGAGGGGTTGGCGGAAAAGGCATGAAAAAACCGGGCGC<br>CAGGCCCGGTTTCCGGTGTCGAAACGCTCTGGCGTCAGGCTTTTGGCAGGGTCACG<br>CCGCGCTGGCCCTGGTATTAAGCTTGGCACTGGCCGTCGTTTTACAACGTC |
| fliC-F | TGCTTTGCTATCGCGACAGTC |
| fliC-R | GGCCATAATCCATCTCCGTCAAA |
| fliC GBlock | TGCTTTGCTATCGCGACAGTCTCCCGGTCACGCGGGTGAAGAACCGTTCCAGGGACT<br>GGCTGGAGGCGATCCGCGCGCACCTGGCCGATGCAGACGCCAACGCCGCGGCCGGC<br>GCGCAGTTGCGCGAGGCGGTGCGCCGCGACTGGATGCTCGAAGGCGCGCATCTCGA<br>GGCATGGGCGGCAGCCTGGCTGCCGGACTGAATCCGGGTTTTTCTCGAACGAGGCC<br>GGTGGCGCAAGCCATCGGCCTTTTTTCATGCCCGGCGTGCCCTGTTGCACGGGAGGG<br>CTAAAGAAAATCGCCGGGGGGTCGATGCAATGGGTGTCGGAACTTCCACCCTCTGC<br>CGGACCAACCGGGGGCGGTTCAGGACCGATATTGGCGAGTCCTCTTCGAAGCATGT<br>AACCCACTGAAGAGGAAGAGAAAAAGAAAATGTTGATTTTTTCTCTAAAGCTCCGC<br>CGGGAAACGCCGATAAACACCATGAACGCGAATTCTTGGGGCACCTGAGCAAGCAG<br>GCCGAGAGATCGCAAGCTCAGGTAACCGAAATAGGTCCTTTGGAGGAAATCACCAT<br>GGCCCGCTAAGCCCGGGAACGGTCACTCACGCGGTACTGGGAGGAAGGGGTGACCC<br>TTCCTCCCTTTTCCCTTTGCGAGGCATGAGAAATGGACGTCGGAAATATCACTTCCC<br>TTTCTACGTTCAAGACCGGCAGGGCCCCGGAGGCCAGCGGCGATATCTTTGCGCCGC<br>GCAGCGCAGCGGATGGCAGCGGCAAACCGTTGCCGGAAGTGACGGCTTCCCGGGAG<br>GCCAGCGAATCTCGCGATGACCTGGGGCTCGCCGTCAGCGACATCCAGTCTTTCGTG<br>CAGAGCGTCAAGCGCAACTTGAACTTCAGCATCGACGACTCCTCTGGCCAGGTCGT<br>GGTCAAGGTGATCGACGGTGACTCGGGCGAAGTCGTGCGGCAAATCCCTTCCGAAG<br>AAGTACTAAAACTTGCCGCCCGGCTCGACGACGTACACAGTGTCCTGTTCGAAACCC<br>GCGCCTGACGCGCGGGTTGAACGACTTGGCATGGTGCTTGCCCTATCGAAGGGATA<br>GGGCAGTACATGCACGGTTTTTTGACGGAGATGGATTATGGCC |

| | |
|---|---|
| pslABC-F | TACGGCCAGTACCTGAACTACCA |
| pslABC-R | TTCCGGCCCGATCTTCATCAA |
| pslABC gBLOCK | GCTTGAATTCGGCAAACGTCCGCCAAGCCCCCCAGGTCGGACCGGCACGCTCGAAC<br>GGCCAACCTGCGTCATGGAACCGCAGGCGCATCCTGCCCAGCCAGCCCGCAGCGCC<br>AGTGGACAGGCGGGTCGTCGAACCGGCGGCTGGCCACCTGGCCGAGCGGCCTGCCC<br>TCACCTTTCGCCCCGCTTCGCTTCCCAGGCCAGAGCGCTCGCGGATTGGCGGCGTCA<br>GATTTCCTCGTCTACTGTTTGGATAAAAGTTTGGCGCCAGAAATACGTCAATAAATT<br>GACTAAAAAAACTTACCCAGACTACGGATATTTCCCTGGGAATGCTAAGATAGCTA<br>TCACAAAGCCACTATCGACGAATGAACCTATTCGACGGGAAAATGACTAAACCGCG<br>TGGCAAATGAAAAATAGTCACTAAATTGACGCTTCACCGCCTTGCTCTTCCCTATCC<br>ACTCAATGGACTGCCCGTGATCGGCAGAGCAAACAACACATCATGACCTACAGGAA<br>GTGCTCCCTCATGAAACGCACCCTCCTCATGCTCGCCATGCTCGCCCTGGCCGCATG<br>CAACACCCCCGCACGGATTCCCGCACCGGACAGCGACACCGTGGACAGCGGCAAGC<br>GTGCCCTGGAAGAACTCGCCAGGCTACCGCCGGCGATGGAGCGGGTGCGCGTCGGG<br>GACACCCTGCGGATCGTCCGCGATGCCGGGGAGATGCCGACCCTCTCGGCGTTCAA<br>CGTCGCCACCATCTATGAACTGACGCTGTACACCGTGCTCAACGACGGCAGCATCTA<br>CTATCCGTTCATCGGTCGCATCCAGGCCGCGCACCGCACGCCGCAGGAAATCGCCA<br>ACGAGCTGACCACCAAGCTCGCGCCGATCTACCGCGAGCCGCGGGTCACGGTGAAC<br>ATCAACCAGGCGCCGGGCAATACGGTGTTCGTCGGCGGCGCGGTGCGCAACCCGTC<br>GGCCGTGCCGATCCCCGCCGCCAACAACATGAAGCTTGCTT |

16

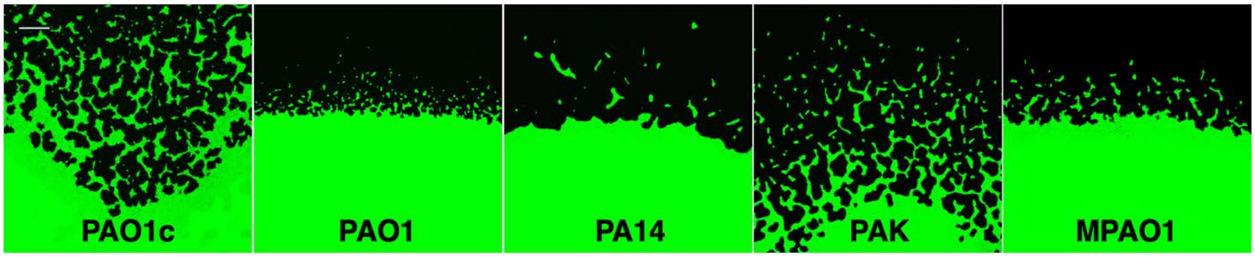

**Fig. S1.** Cluster formation is conserved among wild-type *P. aeruginosa* strains on 0.8% agar. Scale bar represent 100 μm.

**Supplemental Movie Captions:**

Movie S1. *P. aeruginosa* snapping motility. The cluster in the upper right of frame undergoes some cell-cell rearrangement and then snaps to the larger cluster from right-to-left. (5.3 frames per second)

Movie S2. *P. aeruginosa* rapid community contraction. The open area surrounded by cells (spanning 469.87 $\mu m^2$) is covered within 0.76 s.
